# Insights into the Biocontrol Mechanisms of a *Paenibacillus peoriae* Strain on Maize Seedling Blight through Multi-Omics Analysis

**DOI:** 10.1101/2025.04.08.647891

**Authors:** Yue Hu, YiFan Chen, ShengQian Chao, Yin Zhang, LiLi Song, Hui Wang, Yingxiong Hu, BeiBei Lü

## Abstract

Maize seedling blight, caused by the phytopathogenic fungus *Fusarium verticillioides*, is a common and rapidly spreading disease that negatively impacts grain quality and productivity. Control of this pathogen is complicated by its complex infection process and the tendency for resistance to conventional chemical pesticides. The use of biological control agents has been recognized as an environmentally friendly and sustainable solution for the control of plant diseases. In our study, we isolated and identified *Paenibacillus peoriae* 3-B4 from maize leaves, which exhibited an inhibition rate of 59.92% against *F. verticillioides* in greenhouse experiments. Genome sequencing of *P*. *peoriae* 3-B4 revealed a chromosome of 5,912,131 bp, featuring a GC content of 45.51% on average and 5383 annotated coding sequences. Eight gene clusters associated with secondary metabolites with antifungal activity and thirteen genes associated with induced systemic resistance and pattern-triggered immunity were identified. Transcriptomic analysis identified 8,997 differentially expressed maize genes, with key defense genes (e.g., *NPR1*, *bZIP*, *MYB*, *LRR*, *WRKY*) enriched in MAPK signaling and plant–pathogen interactions. 16S rRNA analysis showed significant shifts in microbial communities, particularly with an increase in the abundance of beneficial genera like *Paenibacillus*, *Delftia*, and *Corynebacterium*. The combined analysis of differentially expressed genes and microbial communities indicated that they synergistically enhance pathogen resistance in maize. Our findings outline the potential mechanisms by which *P*. *peoriae* 3-B4 inhibits *F*. *verticillioides* infection and open possibilities of exploiting biological control strategies to control maize seedling blight.

## 1. Introduction

As a highly photosynthetically active C4 crop, maize (*Zea mays* L.) is renowned for its high grain and biomass yields (1), being extensive utilized and playing an integral role in agriculture. This cereal crop is of both great nutritional and industrial importance globally, functioning as a primary source of food for humans and animals alike, and supplying raw materials for a variety of commercial uses (2). Accordingly, major maize disease outbreaks have had a detrimental impact on the economy and the food supply in recent years, posing significant health risks to people and livestock (3). Maize is prone to several fungal diseases, including seedling blight, stalk rot, brown spot, blight, and blackhead disease, all of which severely affect its quality and yield (4,5). Maize seedling blight, with *Fusarium verticillioides* as its causal agent, is among the most common fungal diseases (6). This disease typically occurs during the seedling stage and spreads rapidly, extensively damaging maize plants and resulting in significant yield losses; the disease occurrence could rise to as high as 50% in certain fields (7,8). The main symptoms include withering of leaf edges and gradual yellowing and drying of leaves from the bottom up (9,10). From a physiological point of view, seedling blight impairs photosynthesis, leading to stunting, slow growth, poor development, and leaf wilting or yellowing, adversely affecting the overall growth, weakening maize stalks, and making them more susceptible to lodging (7,11).

*Fusarium verticillioides* (Sacc.) Nirenberg (sinônimo, *Fusarium moniliforme*, Sheldon) is the most widespread fungal pathogen causing seedling blight and rot in various tissues of maize at all growth stages (6,12). In addition to diminishing crop yield and quality, this pathogen could produce fumonisins, which accumulate in maize and pose a risk to human and animal health (13,14). Although traditional chemical control approaches can manage these diseases to some extent, the extensive application of pesticides poses potential risks to the environment, disturbs the ecological equilibrium, and can lead to pathogen resistance (15,16). Comparatively, biological control offers distinct advantages. It is safe for human health, exerts lower environmental impacts than synthetic chemical fungicides, and can achieve sustained, long-term control of pathogens (17,18). Therefore, the development of biocontrol strategy is a treading topic in modern plant disease management (19).

To date, biological management strategies to restrict this fungus include applying microorganisms (such as probiotics, non-toxic fungal strains, and plant growth-promoting rhizobia), antioxidants, and plant extracts, and utilizing genetically engineered disease-resistant crops (20,21). Harnessing microorganisms to manage crop diseases has been increasingly acknowledged as a viable and sustainable alternative to traditional approaches (22). By integrating with plants via, for example, mixing them with roots, seeds, or adding in the soil, biological control agents can effectively target and combat pathogens. Compared to rhizospheric microorganisms, endophytes exhibit superior colonization within plants, evolve in concert with their host plants, and possess inherent colonization advantages (23,24,25). Moreover, endophytic microorganisms reside within plant tissues throughout their life cycle without causing any noticeable harm or adverse effects to the host (26,27). Certain endophytes can have benefits for plants, such as promoting growth and nitrogen fixation and suppressing disease development (28). Microbial fertilizers and biopesticides developed from plant endophytes and their metabolites hold great potential in sustainable agriculture, and have already contributed notably to advancing sustainable maize production (29). As effective biocontrol agents against various plant diseases, several bacterial genera have been proposed (30). For example, the *Paenibacillus* genus, which contains many endophytic bacterial species, has been shown to stimulate plant growth and inhibit pathogen proliferation (31,32). These effects are achieved by inducing plant’s resistance mechanisms or generating inactivating substances (31). In light of these findings, the use of *Paenibacillus* strains and isolated antimicrobial compounds to control plant pathogens may reduce our reliance on chemical fungicides (33).

Although earlier research has revealed that *Paenibacillus* species have the capacity to prevent plant diseases, the mechanisms by which *Paenibacillus peoriae* inhibits phytopathogens have only been reported to a limited extent. In this study, *P*. *peoriae* 3-B4 was identified as an efficacious biological control strain, and its biocontrol mechanism against the phytopathogen *F*. *verticillioides* was studied. Using electron microscopy, we found that the morphological structure of *F*. *verticillioides* was affected by *P*. *peoriae* 3-B4. Genome analysis was carried out to predict the biocontrol potential of *P*. *peoriae* 3-B4. To further elucidate the biocontrol mechanism, transcriptomic analysis and microbial diversity studies, alongside correlation analysis of them, were performed using maize plants upon *P*. *peoriae* 3-B4 induction. Our data provide insights to the inhibitory mechanism of *P*. *peoriae* 3-B4 on plant pathogens and can facilitate develop a novel biocontrol agent, potentially leading to a breakthrough in managing and mitigating fungal pathogen-induced diseases.

## 2. Material and methods

### 2.1 Isolation of endophytic strains and the source of pathogenic strain

The maize plants used in this study were collected from the Zhuanghang Experimental Station of the Shanghai Academy of Agricultural Science in China. Sterilized maize tissue samples, including roots, leaves, ears, and stalks, were sliced into 3mm fragments using surgical scissors and ground in a mortar to obtain a suspension. The suspension was prepared by adding 10 mL of sterilized water per 1 gram of the ground tissue. This suspension was then diluted 10, 10^2^, and 10^3^ times. Following that, 200 μL from each dilution were spread onto three solid media (NB, PDA, and YPD), with 3 replicates for each treatment. The plates were monitored daily for colony and mycelium growth throughout the incubation period. Newly emerged colonies and mycelia were promptly transferred to fresh media for further isolation and purification.

A culture of pathogenic strain *F. verticillioides* 2H12-6 was obtained from the China Specialty Maize Research Centre (Shanghai).

### 2.2 Screening antagonistic endophytes against *F. verticillioides*

To evaluate the antagonistic activities of the isolated strains against fungal pathogen, a bacteria-fungus confrontation assay was performed on PDA. A 5-mm mycelial plug, excised from the edge of a 7-day-old fungal colony on PDA, was positioned in the center of a fresh 90-mm Petri dish. Following a 48-hour incubation period at 37°C and 200 rpm of shaking in Luria-Bertani (LB) broth, the bacterial strains were adjusted to a concentration of 1 × 10^8^ CFU/mL in the suspension. The bacterial suspension was placed in a 5 µL aliquot 25 mm from the fungal plug. As a control, PDA plates with only the fungal mycelial plug were incubated. All plates were incubated at 28°C for 7 days (34). Antifungal activity was determined by evaluating the reduction in radial mycelial growth of the fungal pathogen in the treated samples relative to the control. The percentage inhibition of radial growth (PIRG) was calculated using the formula (PIRG) = ((R1−R2)/R2)×100, where R1 denotes the fungal colony radius on the control plate, and R2 represents the radius in the direction of the antagonist colony (35). This experiment was carried out in three replications.

### 2.3 Identification of the best performing antagonistic endophyte

Taxonomic classification was established by sequencing the 16S rDNA. The 16S rDNA gene fragment and the gyrB gene were PCR-amplified using genomic DNA from *Paenibacillus peoriae* 3-B4 as the template. The primer details can be found in Supplemental Table 8. The amplification products were delivered to Sangon Biotech Co., Ltd. in Shanghai to obtain the sequence. Sequence alignment in NCBI was used to the 16S rDNA and gyrB sequencing findings that were obtained. Eight *Paenibacillus* strains and their corresponding genomes were selected from the NCBI BLAST results, with Scopulibacillus daqui used as an outgroup. The DNA sequence data was analyzed using the BLAST tool. Using MEGA 6.0 software, a phylogenetic tree and a molecular evolutionary analysis were constructed for the data set comprising 16S rDNA sequences.

### 2.4 Scanning Electron Microscope (SEM) Analysis of endophyte-fungal pathogen interaction

To investigate the interaction between the chosen endophytic bacterium and the phytopathogen *Fusarium verticillioides* 2H12-6, 7-day-old mycelial samples were collected from the edge of the inhibition zone. These samples, cultured either on PDA alone or in confrontation with *P. peoriae* 3-B4, were subsequently analyzed using scanning electron microscopy (SEM). The mycelial samples were thinly coated with a 60:40 gold-palladium mixture before being examined under SEM (TM4000PLUS, Hitachi High-Technologies Naka Corporation, Japan).

### 2.5 Greenhouse biocontrol experiments

Greenhouse pot experiments were conducted to evaluate the ability of *P. peoriae* 3-B4 to control maize seedling blight. The inoculum was prepared by culturing *F. verticillioides* on PDA medium for a growth period of between 14 to 21 days until the colonies reached full development. A Petri dish containing spores was submerged in 10 mL of sterile distilled water with 0.0125% Tween® 80. The conidia were removed off the colony surface by gently scraping it with a razor blade. To remove mycelial debris, the spore suspension was filtered through sterile cotton gauze. A hemocytometer was used to determine the spores concentration, and the suspension was made up to 1×10^7^ conidia/mL. The pathogenic fungus *F. verticillioides* 2H12-6 was used to inoculate maize leaves using the needle inoculation method. The experimental group settings are shown in Table 1. Before application, the *P. peoriae* 3-B4 bacterial suspension was diluted 5 times to 5×10^8^ CFU/mL and uniformly sprayed onto the leaf surface. Each treatment group included 10 replicates, with each replicate consisting of 3 maize seedlings. LB liquid medium was applied as the control. Each plant’s disease incidence was noted seven days following inoculation, and the disease index was computed. The 0-5 disease rating scale for maize seedling blight was classified following Zhao et al. (36) . The disease index and control effect were calculated as follows:

**Table 1.**
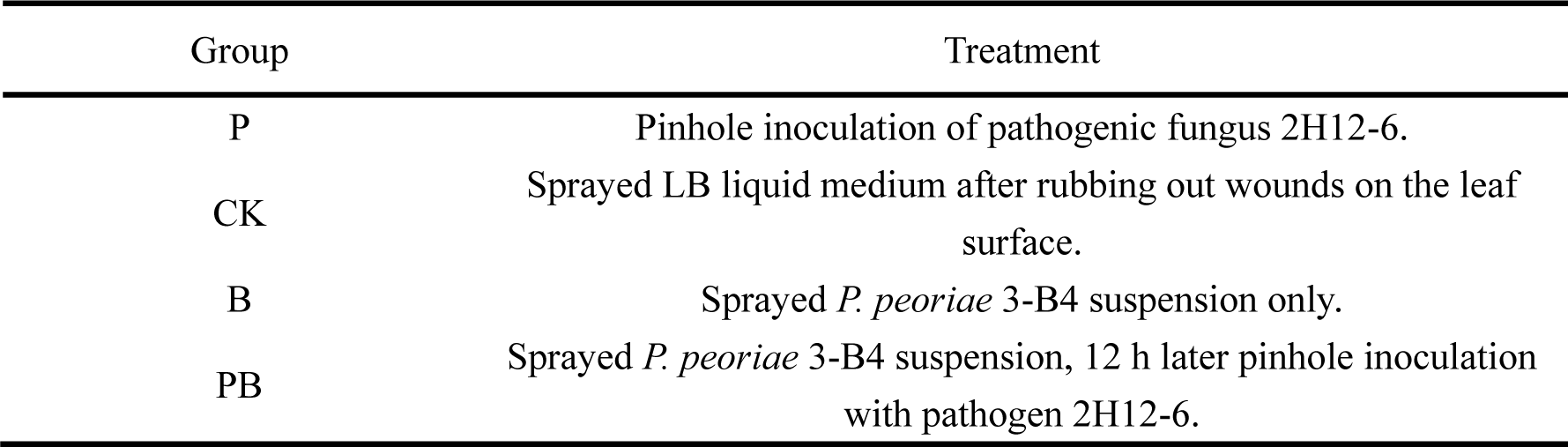
Pot experiment setup.

| Group | Treatment |
| --- | --- |
| P | Pinhole inoculation of pathogenic fungus 2H12-6. |
| CK | Sprayed LB liquid medium after rubbing out wounds on the leaf surface. |
| B | Sprayed *P. peoriae* 3-B4 suspension only. |
| PB | Sprayed *P. peoriae* 3-B4 suspension, 12 h later pinhole inoculation with pathogen 2H12-6. |

Disease index = 100 × ∑ (number of infected maize leaves × relative disease grade) / (total number of maize leaves × highest disease grade).

Control effect (%) = 100 × (disease index of control group − disease index of treatment group) / disease index of control group.

### 2.6 Whole-genome sequencing and analysis of *Paenibacillus peoriae* 3-B4

The *Paenibacillus peoriae* 3-B4 was cultured in LB broth in shaker flasks at 37°C. Genomic DNA was extracted from the cell pellets using the Bacteria DNA Kit (OMEGA) as per the manufacturer’ s protocol, and the purified DNA underwent quality control. DNA quantification was performed using a TBS-380 fluorometer (Turner BioSystems Inc., Sunnyvale, CA). High-quality DNA samples (OD260/280 = 1.8−2.0, >6 µg) were then used for constructing the fragment library.

A minimum of 3 µg of genomic DNA was used for sequencing library construction for Illumina pair-end sequencing of strain 3-B4. Subsequently, the target fragments underwent gel electrophoresis purification, selective enrichment, and PCR amplification. During the PCR step, an index tag was added to the adaptor when necessary, and the library’s quality was assessed afterwards. Finally, the prepared Illumina paired-end library was sequenced using the Illumina NovaSeq 6000 platform (150bp × 2, Shanghai BIOZERON Co., Ltd). The genome sequences of strain *Paenibacillus peoriae* 3-B4 have been uploaded to the NCBI Sequence Read Archive (SRA) Database under the accession number SRR29484736.

For the prokaryotic strain 3-B4, gene models were generated using an ab initio prediction method. GeneMark was employed for this purpose, and the resulting models were subjected to blastp analysis against the NCBI non-redundant (NR) database, KEGG, and COG for functional annotation.

### 2.7 Transcriptome sequencing and data analysis

Gene expression profiles were assessed for Group B (treated with *P. peoriae* 3-B4 suspension on maize leaves) and Group CK (untreated control) in pot experiments. Five biological replicates were performed, and there were 10 samples totally.

Total RNA was extracted from maize leaves using Trizol Reagent (Invitrogen Life Technologies), followed by the assessment of concentration, quality, and integrity with a NanoDrop spectrophotometer (Thermo Scientific). Three micrograms of RNA were utilized as input for RNA sample preparation. The final products were quantified using an Agilent high-sensitivity DNA assay on a Bioanalyzer 2100 system (Agilent) following purification with the AMPure XP system. The sequencing library was sequenced by Ltd. using the Illumina NovaSeq 6000 platform. Following standardisation of the expression data using FPKM, the read count values for each gene were compared to the gene’s original expression using HTSeq(0.9.1) statistics. Subsequently, the DESeq (1.30.0) software was employed to evaluate the differences in gene expression, utilising the following screening settings: P value < 0.05 indicates a substantial expression differential multiple |log2FoldChange| > 1.

The complete set of genes was aligned with the terms utilized in the Gene Ontology database, thereby establishing the total number of differentially enriched genes attributed to each term. A GO enrichment analysis of the differentially expressed genes was conducted using the topGO package, with p-values calculated through the hypergeometric distribution method (significance threshold set at p-value < 0.05). This enabled the identification of GO terms with significantly enriched differentially expressed genes, thereby highlighting the primary biological functions associated with these genes. Enrichment analysis of KEGG pathways for the differentially expressed genes was conducted using ClusterProfiler (v3.4.4), with emphasis on pathways exhibiting significant enrichment and a p-value of less than 0.05.

The raw data were deposited in the NCBI Sequence Read Archive (SRA) Database under the accession number SRR29873971, SRR29873972, SRR29873973, SRR29873974, SRR29873975, SRR29873976, SRR29873977, SRR29873978, SRR29873979, SRR29873980.

### 2.8 16S rRNA sequencing and bioinformatic analysis

DNA was extracted from maize leaves in the CK group (5 control samples) and the B group (5 samples treated with biocontrol bacteria) using the OMEGA Soil DNA Kit (M5635-02, Omega Bio-Tek, Norcross, GA, USA). PCR was utilised to amplify the V3–V4 region of the bacterial 16S rRNA genes. Vazyme VAHTSTM DNA Clean Beads (Vazyme, Nanjing, China) were employed for the purification of PCR amplicons, while the Quant-iT PicoGreen dsDNA Assay Kit (Invitrogen, Carlsbad, CA, USA) was used for the quantification of results. Subsequently, the amplicons were pooled in equal quantities and subjected to pair-end 2×250 bp sequencing at the Shanghai Personal Biotechnology Co., Ltd. (Shanghai, China) facility, utilising the Illumina NovaSeq platform and the NovaSeq 6000 SP Reagent Kit (500 cycles).

The microbiome bioinformatics analysis was performed using QIIME2 2019.4. QIIME2 and the R package (v3.2.0) were mostly used to analyze sequence data. Alpha diversity indices at the ASV level, including the Chao1 richness estimator, Shannon diversity index, and Simpson index, were computed based on the ASV table in QIIME2 and presented as box plots. All the raw reads have been deposited under the BioProject accession number PRJNA1134884 in the NCBI BioProject database.

### 2.9 Quantitative real time-PCR (qRT-PCR) analysis

To confirm the transcriptome sequencing results, a random selection of eleven DEGs associated with plant disease resistance genes was subjected to quantitative real-time PCR (qPCR) analysis. The specific primers used for this analysis are provided in Supplemental Table 9, and the expression trends were compared to those observed in the transcriptomic data. Total RNA was extracted from maize leaves by Trizol Reagent (Invitrogen Life Technologies). The same samples used for RNA-seq were also employed for qRT-PCR analysis. RNA was subsequently reverse-transcribed into cDNA utilizing the PrimeScript TM 1st Strand cDNA Synthesis Kit. This cDNA served as the template for qPCR. Subsequently, the calculated Ct values were employed in the determination of the relative expression of each gene through the 2^−ΔΔCt^ method, with β-actin as the housekeeping gene. Each sample was analyzed in at least three technical replicates.

### 2.10 Data Analysis

Each assay was conducted in at least three independent experiments. All values are expressed as mean ± standard error based on a minimum of three replicates. Variance was analyzed using ANOVA in GraphPad Prism 8 (v8.0.1.244).

## 2. Results

### 2.1 Isolation, screening, and classification of antagonistic endophytic strains against *F*. *verticillioides*

A total of 147 bacterial strains and one actinomycete strain were isolated from *Zea mays L.* in this study. Genomic DNA was isolated from these strains, followed by amplification of the 16S rDNA fragment. Species classification was completed after sequencing (Supplemental Table 1). The bacteria were numbered, preserved, and recorded for further analysis. Colony morphological characteristics are shown in Figure 1A and Supplemental Table 2. Statistical analysis of these strains revealed that *Bacillus*, *Pantoea*, *Enterobacter*, *Paenibacillus,* and *Brevibacillus* accounted for 45, 8, 7, 4, and 4% of the total isolates, respectively. The isolated strains belong to 26 genera from 16 families. Among these, high abundance genera were *Bacillus*, *Enterobacter*, and *Pantoea* (Figure 1B), suggesting that these three genera were dominant in the culturable endophytic flora of maize tissues.

**Figure 1.**
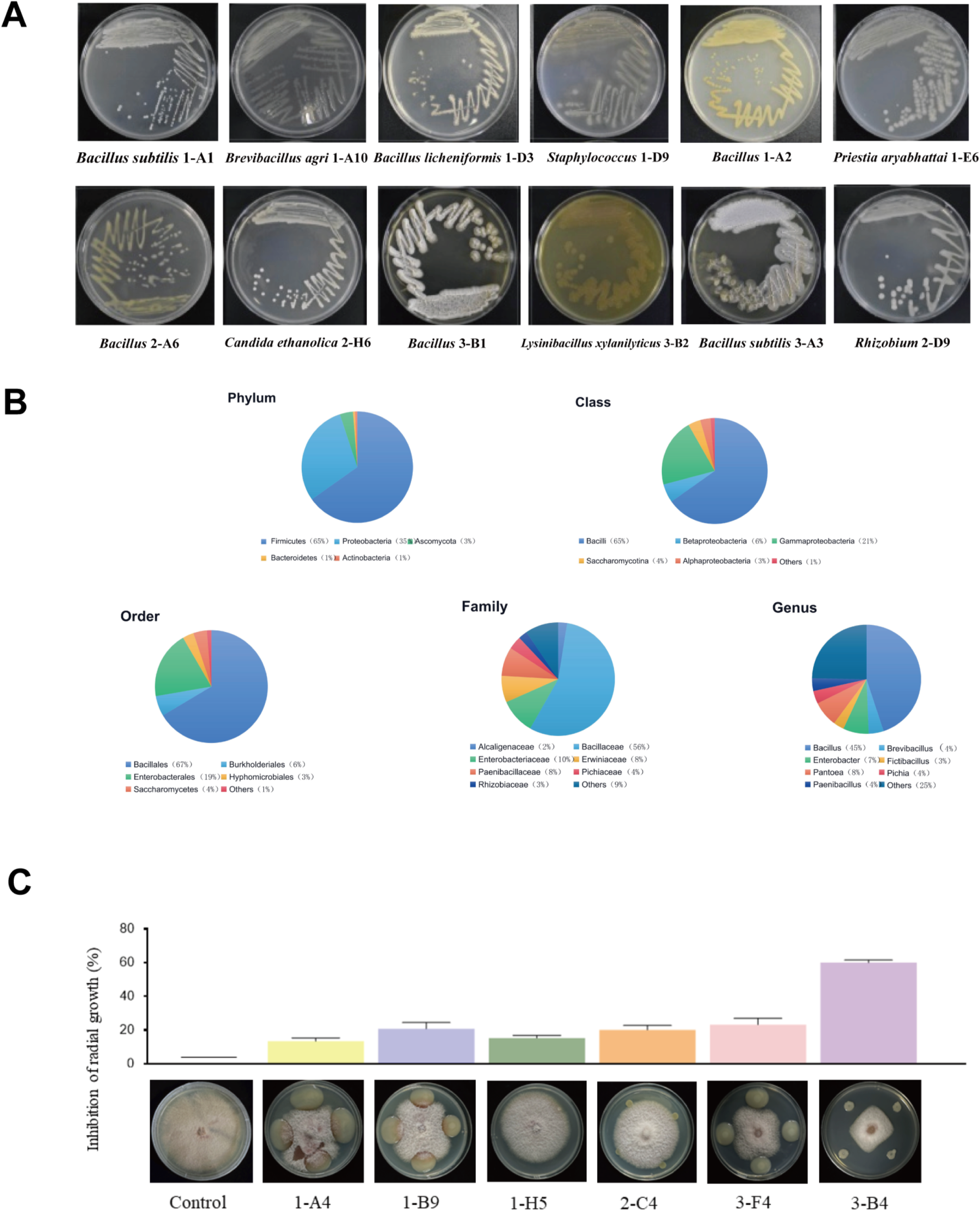
Isolation, identification, and antagonistic screening of endophytic bacteria from maize. **(A)** Colony morphology of isolated bacteria. **(B)** Distribution and diversity analysis of 148 maize endophytic bacterial strains at different taxonomic levels. **(C)** Fungal inhibition assays of certain strains against *Fusarium verticillioides* 2H12-6.

The antifungal effects of the isolated endophytic bacteria were verified through plate confrontation assays. Strain 3-B4 exhibited the greatest effective inhibition on *F*. *verticillioides* 2H12-6 growth, as indicated by the smallest growth area of fungal colonies (32.9 mm in diameter), and the largest inhibition rate (59.92%) (Supplemental Table 3). In contrast, the bacterial growth areas of strains 1-A4, 1-B9, and 3-F4 were significantly larger than those of strain 3-B4, and the fungal mycelium was not noticeably inhibited (Figure 1C).

### 3.2 Morphological and molecular identification of *P*. *peoriae* 3-B4

The microscopic image showed that the colonies of strain 3-B4 were yellow in color, with intact margins. Upon Gram staining, the colonies exhibited a reddish purple coloration, confirming that strain 3-B4 was a Gram-negative bacterium (Supplemental Figure 1B, C).

The BLAST search and sequence comparison using the 16S rDNA sequence of strain 3-B4 in the GenBank nucleotide database revealed a high sequence similarity to *P. peoriae*, indicating a close phylogenetic relationship with other *Paenibacillus* species. The molecular identification aligned with our morphological and physiochemical characterization, confirming that strain 3-B4 belongs to the species *P. peoriae*. (Supplemental Figure 1A).

### 3.3 Inhibition of the morphological structure of *F*. *verticillioides* by *P*. *peoriae* 3-B4

The antifungal activity of *P*. *peoriae* 3-B4, which produces fusaricidin with potent broad-spectrum activity, was studied to assess its impact on the fungal pathogen structure. Morphological observations using electron microscopy revealed that during the infection process, *F*. *verticillioides* 2H12-6 exhibited tumor-like swollen structures in pure cultures, representing specific morphological changes occurring (Figure 2A, B, C). In contrast, during the antagonistic interaction between *F*. *verticillioides* 2H12-6 and *P*. *peoriae* 3-B4, the tumor-like structures were not observable, the number of spores decreased, and the hyphae showed breakage (Figure 2D, E, F). This suggests that *P*. *peoriae* 3-B4 may inhibit the infection structures of *F*. *verticillioides* hyphae, thereby further controlling pathogen growth.

**Figure 2.**
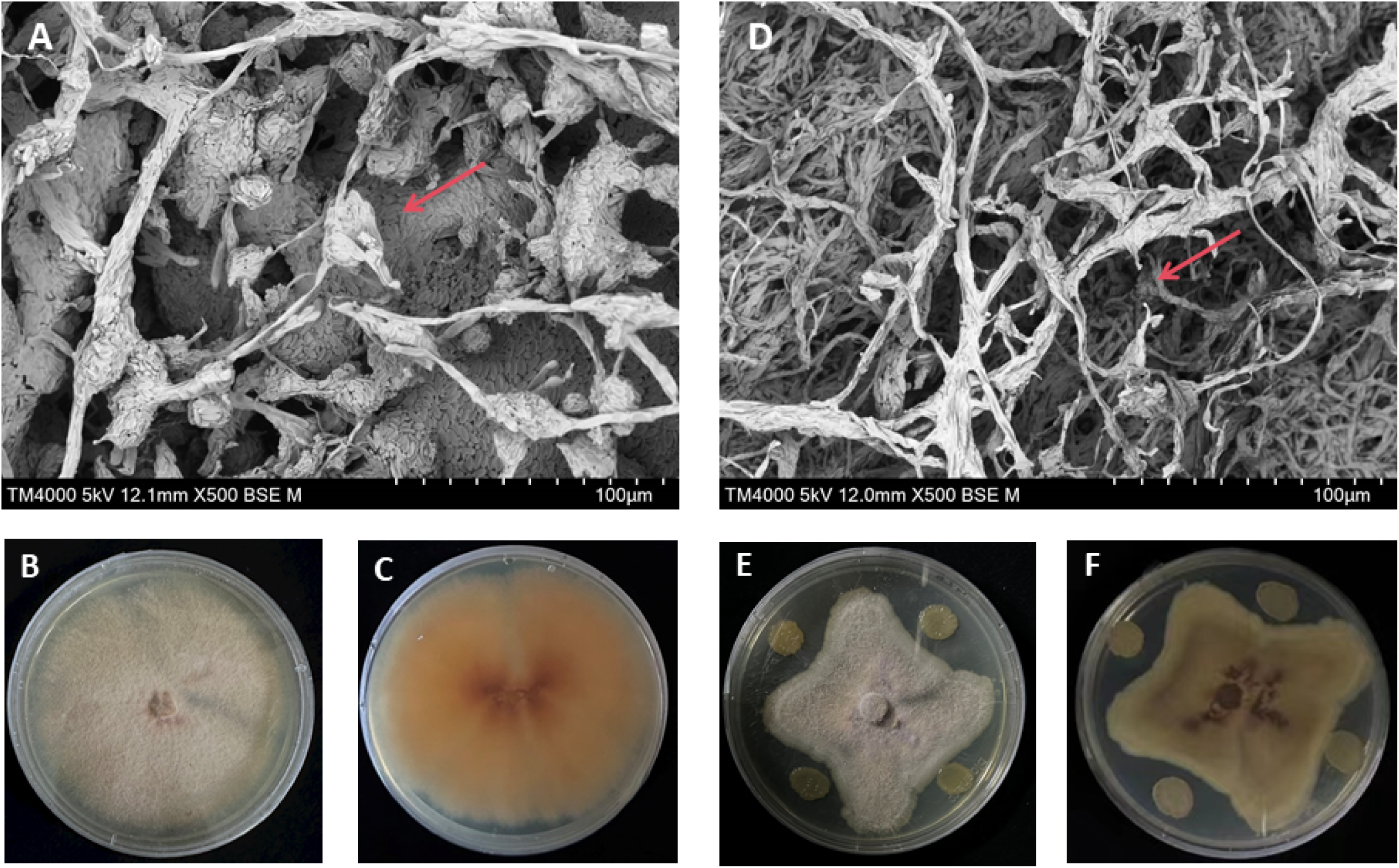
Morphological alterations of *Fusarium verticillioides*. (A, B, **C)** *Fusarium verticillioides* 2H12-6 in single-strain culture. **(D, E, F)** *Fusarium verticillioides* 2H12-6 cultured with *Paenibacillus peoriae* 3-B4.

### 3.4 Biocontrol efficacy of *P*. *peoriae* 3-B4 against maize seedling blight

The efficacy of *P*. *peoriae* 3-B4 as a biocontrol agent against maize seedling blight was validated through pot experiments. On plants inoculated only with fungal pathogen *F*. *verticillioides* (Group P), leaf spots rapidly expanded, leaves curled and shriveled, and the disease extended to the whole seedling, which dried up and died (Figure 3B). In contrast, when the biocontrol bacterium was used to pretreat the maize leaves (Group PB), reduced disease symptoms were observed, with suppressed spots and no visible leaf yellowing (Figure 3D). The growth of plants sprayed only with the biocontrol bacterium (Group B), as shown in Figure 3C, was comparable to the control group (Group CK), as shown in Figure 3A, indicating that plant growth wasn’t affected by the biocontrol bacterium’s application. Seven days after inoculation with *F. verticillioides*, the morbidity of maize seedlings in group P exceeded 80%. At this point, the disease condition index was calculated. The pre-treatment group (Group PB) that received the biocontrol bacterium exhibited a significant decrease in the occurrence and severity of maize leaf disease and in maize seedling blight index compared to Group CK. The biocontrol effect was 77.23%, and the disease index was reduced by 59.78% at seven days after inoculation with *F*. *verticillioides* (Table 2).

**Figure 3.**
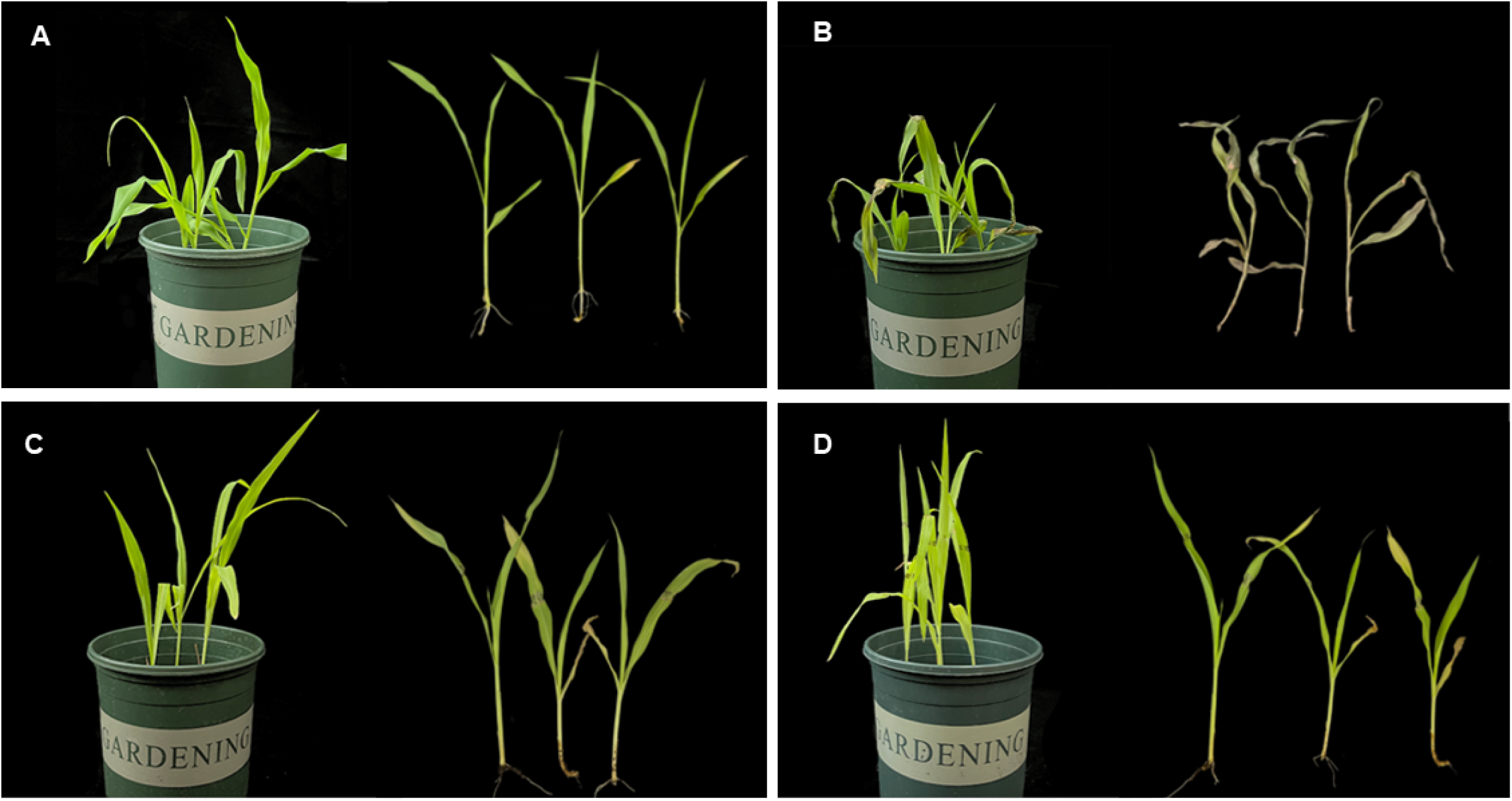
Biocontrol efficacy of *Paenibacillus peoriae* 3-B4 in maize seedlings. **(A)** Maize seedlings sprayed with LB liquid medium. **(B)** Maize seedlings inoculated with pathogenic fungus *Fusarium verticillioides* 2H12-6. **(C)** Maize seedlings treated with biocontrol bacterium *P. peoriae* 3-B4. **(D)** Maize seedlings inoculated with pathogenic fungi after pretreatment with biocontrol bacterium *P. peoriae* 3-B4.

**Table 2.** Greenhouse disease control assays of biocontrol bacterium *Paenibacillus peoriae* 3-B4 against maize seedling blight.

| Group | Disease index (%) | Biocontrol efficacy (%) |
| --- | --- | --- |
| CK | 0 | - |
| B | 0 | - |
| P | 80.34 | - |
| PB | 20.56 | 77.23 |

### 3.5 General biological and genomic characteristics of *P*. *peoriae* 3-B4

*P. peoriae* 3-B4’s whole genome comprises a circular DNA structure with 5,839,239 base pairs and a mean G+C content of 45.51%. An overview of the assembly details and genomic characteristics is presented in Figure 4E and Supplementary Tables 4 and 5. The genome revealed 5383 predicted genes, including 17 ribosomal RNA operons and 79 tRNAs (Supplementary Table 6).

**Figure 4.**
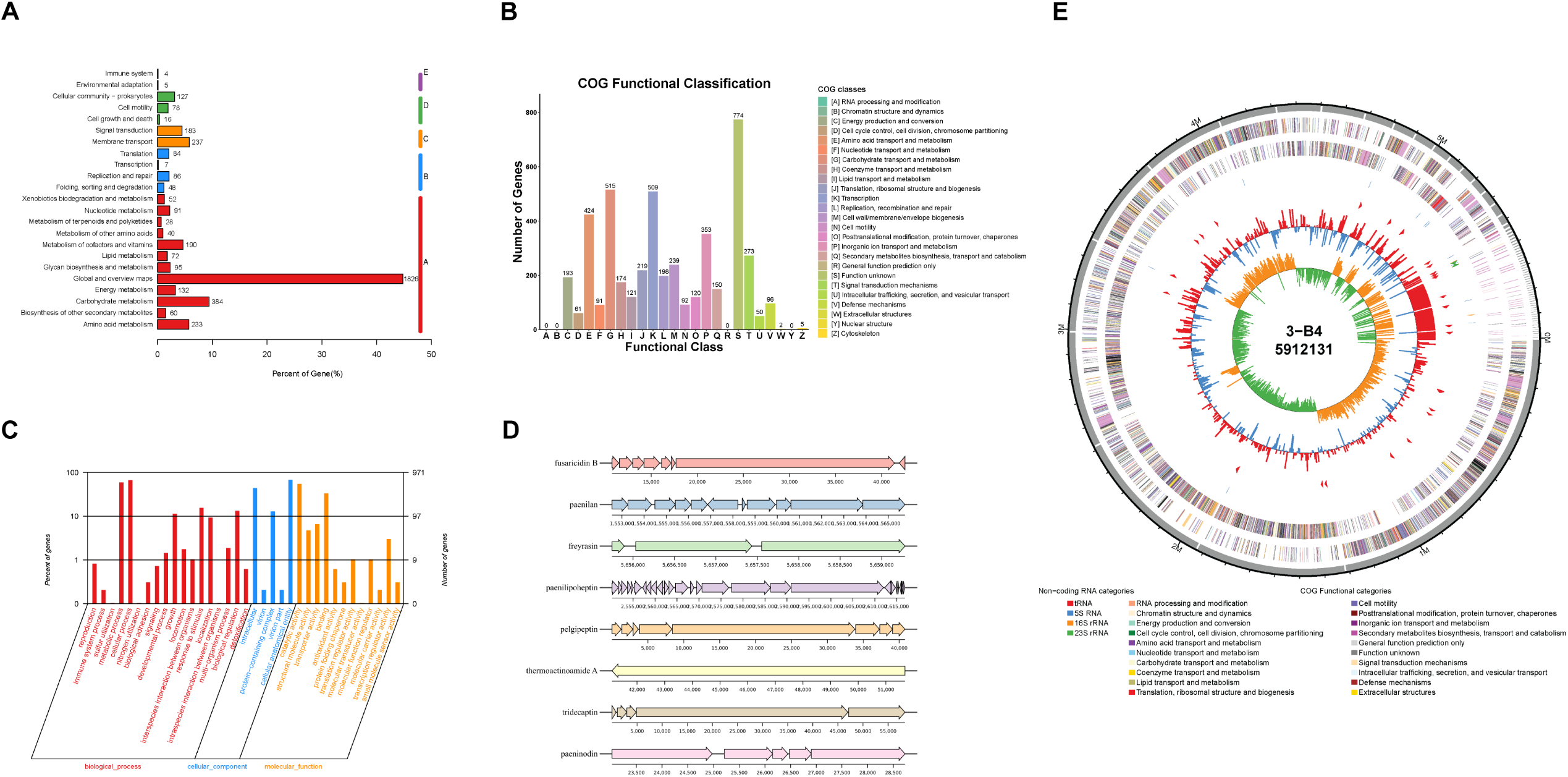
Whole genome analysis of *Paenibacillus peoriae* 3-B4. **(A)** KEGG functional annotation classification statistics. **(B)** COG functional annotation classification statistics. **(C)** GO functional annotation classification statistics. **(D)** Cluster of secondary metabolite-related genes annotated by antiSMASH. Each band represents a gene, and the color and shape indicate the functional category or expression of the gene. Genomic locations indicated by numerical labels in the figure. The direction of gene transcription is indicated by an arrow. **(E)** Graphical circular map of the *P. peoriae* 3-B4 genome.

A total of 2731 functional genes were identified and mapped using the Kyoto Encyclopedia of Genes and Genomes (KEGG) database. These genes were categorized into five groups of KEGG metabolic pathways. The largest share of annotated genes, amounting to 1826, was linked to cellular process metabolic pathways. Among these, most were concentrated in Global and overview maps, Carbohydrate metabolism, and Amino acid metabolism (Figure 4A, Supplementary Figure 2).

Gene function annotation of *P*. *peoriae* 3-B4 using the Gene Ontology (GO) database is shown in Figure 4C. Among the three GO categories, Biological process contained the largest collection of enriched genes, particularly metabolic and cellular processes. This category was followed by Molecular function, where catalytic activity and binding-related genes were the most abundant, while genes related to Molecular transducer activity were the least represented. Figure 4B presents the Clusters of Orthologous Groups of Proteins (COG) annotation analysis for *P. peoriae* 3-B4. Of all coding genes, 4229 were annotated to the COG database, accounting for approximately 78.56%. These genes were distributed in 24 homologous gene clusters. After removing 774 genes with unknown functions, three major homologous gene clusters with a large number of predicted genes were obtained: carbohydrate transport and metabolic functions (515 genes, 12.18%), transcriptional functions (509 genes, 12.03%), and amino acid transport and metabolic functions (424 genes, 10.03%).

### 3.6 Genes/gene clusters involved in antibiotic synthesis and plant resistance induction

*Paenibacillus* species have been found to exhibit potent broad-spectrum antifungal activity against various fungi, including *Fusarium oxysporum*, *Fusarium verticillioides*, and *Fusarium pseudograminearum*, among others (37). In the genome sequence of *P. peoriae* 3-B4, genes associated with secondary metabolism were identified in the genome sequence of *P. peoriae* 3-B4 using the AntiSMASH database to explore potential secondary metabolites linked to antifungal activity. Gene clusters involved in several pathways of biosynthesis were identified, four of which encode non-ribosomal peptide synthetases (NRPSs). Notably, fusaricidin B, paenilan, freyrasin, paenilipoheptin, pelgipeptin, paeninodin, thermoactinoamide A, and tridecaptin exhibited significant antifungal activity against fungal pathogens (Figure 4D).

The *P. peoriae* 3-B4 genome was analyzed using KEGG to identify key genes involved in induced systemic resistance (ISR). These included 10 ISR-related genes and 3 genes linked to pattern-triggered immunity (PTI), all of which were analyzed for their roles in resistance inducer biosynthesis. The characteristics of these genes in *P. peoriae* IBSD35, GXUN15128, and HS311 were compared using the NCBI database (Table 3). The results indicated that the genomes of these four *P. peoriae* strains contained key genes related to volatile organic compounds in ISR, including those encoding the production of 2,3-butanediol, methanethiol, isoprene, and peptidoglycan, with their sequences in the *P. peoriae* 3-B4 genome showing over 95% homology to the corresponding sequences in strains IBSD35, GXUN15128, and HS311.

**Table 3.** Comparative analysis of genes related to the synthesis of resistance inducer strains.

| Gene name | Plant resistance type | Resistance inducers | Location | Identity (%) |  |  |
| --- | --- | --- | --- | --- | --- | --- |
|  |  |  |  | *P.peoriae*<br>.IBSD35 | *P.peoriae*.<br>GXUN15128 | *P.peoriae*.<br>HS311 |
| alsD | ISR | 2,3-Butanedio | 1,291,944–1,291,198 | 99.6 | 99.6 | 97.58 |
| bdhA | ISR | 2,3-Butanedio | 425,759–426,820 | 99.43 | 99.71 | 99.71 |
| alsS | ISR | 2,3-Butanedio | 1,293,804–1,292,107 | 96.09 | 96.28 | 96.28 |
| alsR | ISR | 2,3-Butanedio | 912,647–911,748 | 99.67 | 97.99 | 97.66 |
| ilvN | ISR | 2,3-Butanedio | 541,932–542,417 | 100 | 100 | 100 |
| ispE | ISR | Isoprene | 18,527–19,390 | 99.3 | 99.65 | 98.94 |
| ispF | ISR | Isoprene | 50,295–49,819 | 96.2 | 95.57 | 96.2 |
| metE | ISR | Methanethio | 310,364–312,682 | 98.45 | 98.58 | 98.58 |
| metH | ISR | Methanethio | 327,895–324,515 | 99.83 | 99.74 | 99.91 |
| mmuM | ISR | Methanethio | 535,045–534,098 | 98.41 | 99.37 | 98.73 |
| tuf | PTI | Elongation factor Tu | 27,186–25,996 | 98.99 | 100 | 100 |
| dacA | PTI | Peptidoglycan | 34,599–35,555 | 100 | 100 | 100 |
| dacA | PTI | Peptidoglycan | 63,246–61,858 | 99.78 | 98.48 | 98.48 |

### 3.7 Transcriptomic profiles of *Zea may* L. upon *P*. *peoriae* 3-B4 application

We constructed a total of 10 libraries for CK and B, with raw reads of 3.70–4.97 Gb and approximately 3.60–4.83 Gb of clean reads. The values for Q20 and Q30 were above 96.19% and 93.35% (Supplemental Table 7). Comparison efficiency is defined as the percentage of mapped reads to clean reads. In our mapping process, the comparison efficiency was greater than 70% and these mapped reads were fully assembled, showing a high-matching rate to the reference genome and no signs of contamination. The ratio efficiency is over 88.92%, and more than 96.50% of reads were located in exonic regions. The unigenes annotated from the NR database were aligned with various species, with 95.02% of them identified as belong to maize, which is generally consistent with previous reports. Overall, these results demonstrate that the sequencing was of high quality and met the necessary criteria for further analysis.

In our differential expression analysis, 8997 differentially expressed genes (DEGs) were detected when comparing CK with B. Among these, 5059 showed upregulated, while 3938 showed downregulated (Figure 5A), indicating significant changes in maize gene expression after *P*. *peoriae* 3-B4 application. The volcano plot in Figure 5C illustrates the statistical significance and fold-change of gene expression differences.

**Figure 5.**
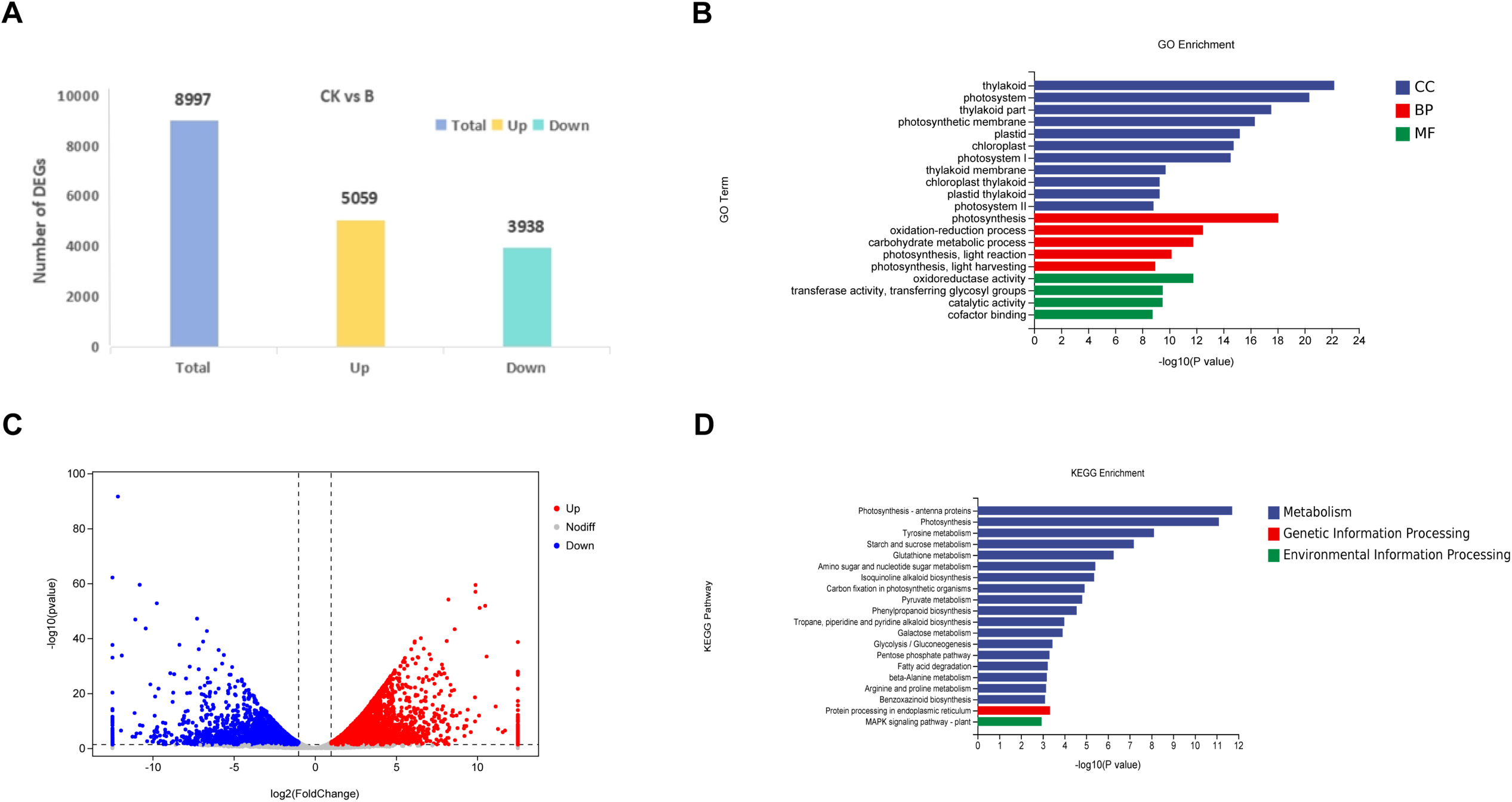
Transcriptional level analysis of *Paenibacillus peoriae* 3-B4 on maize leaves. **(A)** Analysis of differentially expressed genes between CK and B groups. **(B)** GO enrichment plot of differentially expressed genes. **(C)** Volcano map of differentially expressed genes. **(D)** KEGG enrichment plot of differentially expressed genes.

Eleven genes were randomly chosen from the identified DEGs and analysed by quantitative PCR (qRT-PCR) to confirm the RNA sequencing (RNA-Seq) results. β-actin served as a control gene. The qPCR results showed a similar expression pattern to the transcriptome data’s expression count values for each gene, thereby validating the RNA-Seq-generated expression results (Supplementary Figure 3).

#### 3.7.1 GO functional enrichment analysis of the DEGs

GO enrichment analysis of DEGs identified 4 molecular function terms, 5 biological process terms, and 11 cellular component terms (Figure 5B). Within cellular components, the most significantly enriched terms were thylakoid membrane system-related categories, including ‘thylakoid’, ‘photosystem’, and ‘chloroplast thylakoid’, collectively representing 38% of cellular component DEGs. These were followed by ‘plastid thylakoid membrane’ and ‘photosynthetic membrane’ , which we clarify both refer to specialized membrane structures within chloroplasts. Notably, the term ‘plastid’ in this context specifically refers to chloroplast subcompartments rather than other plastid types. For molecular functions, dominant categories included ‘oxidoreductase activity’ and ‘transferase activity’ , particularly those transferring glycosyl groups. Biological process DEGs were predominantly associated with photosynthetic pathways, including ‘photosynthesis, light reaction’ and ‘light harvesting’. This comprehensive analysis indicates that *P. peoriae* 3-B4 primarily influences maize leaf metabolism through modulation of photosynthetic machinery, redox homeostasis, and carbohydrate metabolism pathways.

#### 3.7.2 Biological pathways mediating the effects of *P*. *peoriae* 3-B4

To better understand the biological pathways through which strain 3-B4 affects maize, KEGG functional annotation analysis was conducted on all DEGs identified by RNA-Seq. The results revealed that 8997 DEGs showed enrichment in 112 KEGG pathways (Figure 5D). In these pathways, DEGs were most significantly enriched in protein processing in the endoplasmic reticulum and plant–pathogen interactions (Table 4). According to the KEGG annotation results, we selected the following five KEGG pathways with significantly enriched DEGs for subsequent analysis: photosynthesis-antenna protein (zma00196), mitogen-activated protein kinase (MAPK) signaling pathway—plant (zma04016), ABC transporters (zma02010), protein processing in endoplasmic reticulum (zma04075), and plant–pathogen interaction (zma04626).

**Table 4.** KEGG pathway enrichment of DEGs.

| Pathway ID | Pathway | DEGs | P-value | Adjust P-value |
| --- | --- | --- | --- | --- |
| zma04075 | Protein processing in endoplasmic reticulum | 104 | 0.003728596 | 0.017123921 |
| zma04626 | Plant-pathogen interaction | 85 | 0.025746243 | 0.086284707 |
| zma00520 | Amino sugar and nucleotide sugar metabolism | 79 | 1.17212E–05 | 0.000215533 |
| zma00500 | Starch and sucrose metabolism | 76 | 6.21356E–06 | 0.000154096 |
| zma00940 | Phenylpropanoid biosynthesis | 74 | 0.003291745 | 0.015699092 |
| zma04016 | MAPK signaling pathway - plant | 67 | 0.004918376 | 0.021030297 |
| zma00010 | Glycolysis / Gluconeogenesis | 66 | 0.001364274 | 0.010565242 |
| zma00195 | Photosynthesis | 59 | 6.00827E–16 | 7.45025E–14 |
| zma00480 | Glutathione metabolism | 57 | 2.15614E–06 | 6.68402E–05 |
| zma00564 | Glycerophospholipid metabolism | 55 | 0.001973083 | 0.012017907 |
| zma00620 | Pyruvate metabolism | 54 | 1.21672E–05 | 0.000215533 |
| zma00710 | Carbon fixation in photosynthetic organisms | 49 | 3.22756E–05 | 0.000444686 |
| zma00230 | Purine metabolism | 47 | 0.019766614 | 0.072090005 |
| zma00620 | Pyrimidine metabolism | 41 | 0.000431791 | 0.004662085 |
| zma00710 | Glyoxylate and dicarboxylate metabolism | 39 | 0.010652104 | 0.044028696 |

In the photosynthesis-antenna protein pathway, the expression levels of 27 DEGs exhibited a notable increase in the biocontrol-treated leaves (Group B) relative to the control leaves (Group CK) (Figure 6A). Among these DEGs, 5 *Lhca* genes, 5 *Lhcb* genes, and 7 *Cab* genes were upregulated, while 1 *LHBC* gene was downregulated. The capture and transfer of light energy is the main function of these genes. The *Lhca* gene family encodes Light-harvesting Complex I (LHCI) proteins, a series of pigment-binding membrane proteins in Plant Photosystem I (PSI). In total, 67 DEGs were identified as being associated with the MAPK signaling pathway, with 25 showing increased expression and 42 showing decreased expression (Figure 6B). Notably, the MPK3, SAPK4, Rbohe, and CAT2 genes exhibited increased expression, while the MPK1, MPK5, MPK12, WRKY24, and PYL4 genes exhibited decreased expression. Among the MKK5 genes, two had increased expression, and one had decreased expression, indicating tha their different functions in MAPK signaling pathway. In the plant-pathogen interaction pathway, 85 DEGs were identified, with 40 upregulated and 45 downregulated. Differential expression was identified for calcium-binding protein (CML)-encoding genes, PTI1-like tyrosine-protein kinase (PTI) genes, 3-ketoacyl-CoA synthase (KCS), and Calcium-dependent protein kinase (CPK) genes. Furthermore, decreased expression was observed for five Plant Immune Receptor (RPM1) genes and five WRKY transcription factors. The functionality of these genes is of vital importance in plant responses to various environmental stresses. RPM1 is involved in recognizing pathogen effector-induced modifications and triggering immune responses. WRKY transcription factors, on the other hand, act as regulators of gene expression, modulating the transcription of genes involved in stress responses and defence mechanisms. In protein processing in the endoplasmic reticulum pathway, 104 DEGs were identified, with 41 exhibiting increased expression and 63 showing decreased expression (Supplementary Figure 4). Among these, eight DnaJ/40 heat shock proteins demonst rated increased expression. Additionally, five HSP70 genes exhibited increased expression, and they contribute to protein folding and the stress response; whereas four PDI genes showed decreased expression, and these genes are associated with protein folding and the maintenance of redox homeostasis.

**Figure 6.**
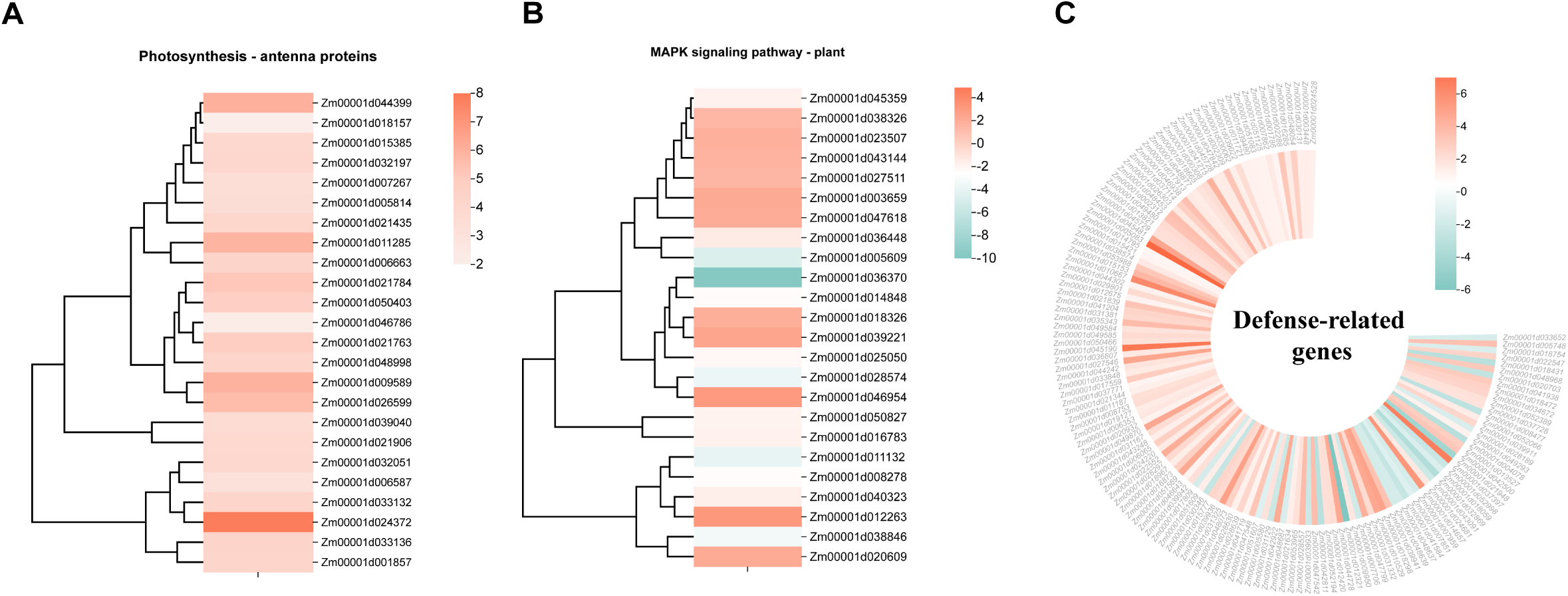
Heatmap of expression levels of potential genes involved in plant disease resistance. The red color represents upregulation, and the blue represents downregulation. (A) Genes involved in photosynthesis-antenna protein. (B) Genes involved in MAPK signaling pathway—plant. (C) Defense-related genes.

#### 3.7.3 Potential functional genes and transcription factors related to pathogen resistance

In our transcriptome analysis, 144 candidate genes, including 14 KINs, 5 LECRKs, 3 RLKs, 2 RLPs, 1 CN, 1 CNL, 1 LEC, and 1 TRAN, that may be related to resistance against maize seedling blight were identified. These genes were mainly related to 7 RPP13-like proteins, 35 HSP, 1 HT1, 23 bHLH, 8 WAK, 8 GSTs, 1 LIPOXYGENASE, 1 remorin, 1 pan1, 1 WRKY, 6 bZIP, 27 LRR, 3 FBL, and 3 CCT. Of these, bHLH, bZIP, and WRKY are transcription factors. These genes are presented in Figure 6C and Table 5. The gene Zm00001d042111, which encodes Nonexpressor of Pathogenesis-Related Genes 1 (NPR1), is a key regulator in the salicylic acid (SA) signaling pathway (38). Its expression was significantly upregulated, showing a 3.69-fold increase. NPR1, a regulatory protein integral to plant immune responses, is essential for the expression of SA-mediated pathogenesis-related (PR) genes (39,40). NPR1 represents a pivotal node in the interconnection between SA signaling and PR gene expression and is of significant importance for plants to acquire systemic acquired resistance (41,42).

**Table 5.**
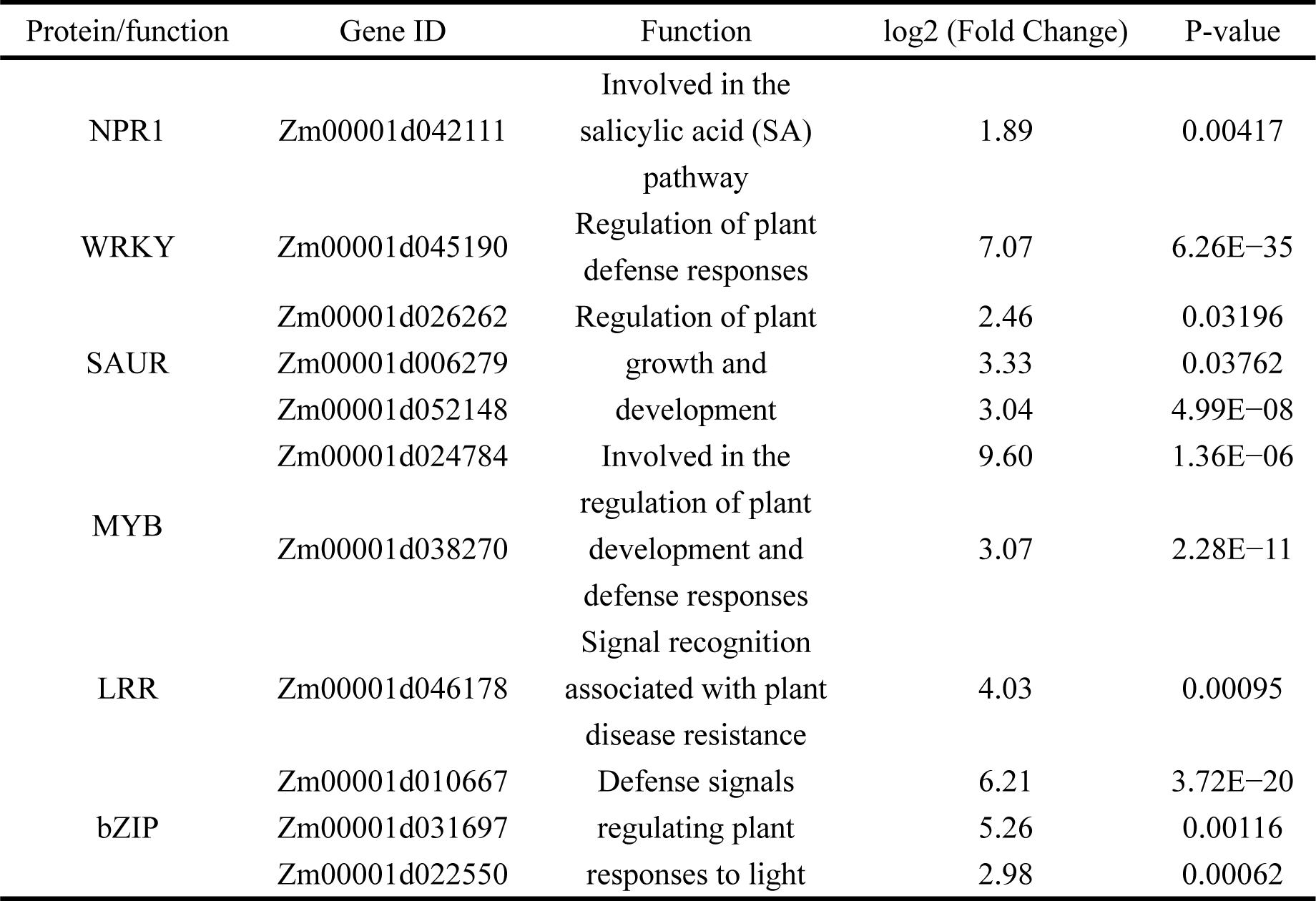
Genes related to plant hormone signal transduction and disease resistance.

### 3.8 Shifts in the microbial community composition of maize leaves treated with the biocontrol bacterium *P*. *peoriae* 3-B4

Our transcriptomic analysis revealed that the plant–pathogen interaction pathway showed the most significant divergence among differentially expressed genes (DEGs). Therefore, this could be caused by phyllospheric microorganisms. Thus, we sequenced 16 leaves of both groups, compared the bacterial species and abundance in CK and B leaves, and screened the candidate microorganisms involved in plant disease resistance. We obtained 133,801 and 137,725 effective sequence reads by 16S rRNA diversity sequencing. A total of 1,372 amplicon sequence variants (ASVs) were identified, with 66 ASVs shared between the two groups.

Microbial composition analysis revealed that the 5 most predominant phyla in the control group were Proteobacteria (67.60%), Firmicutes (28.01%), Actinobacteriota (2.08%), Bdellovibrionota (2.01%), and Acidobacteriota (0.19%). In contrast, the biocontrol treatment group displayed a different pattern, with the top 5 phyla being Proteobacteria (41.23%), Firmicutes (32.87%), Actinobacteriota (20.51%), Bacteroidota (4.71%), and Myxococcota (0.54%). *Limnobacter* (17.92%), *Escherichia-Shigella* (5.46%), *Sphingomonas* (4.14%), *Enterococcus* (2.87%), and *Staphylococcus* (2.68%) were the top five genera in the control group, while *Corynebacterium* (19.96%), *Paenibacillus* (17.29%), *Delftia* (13.26%), *Escherichia-Shigella* (10.60%), and *Chryseobacterium* (4.71%) dominated the biocontrol treatment group (Figure 7A). The α-diversity analysis revealed a statistically significant reduction in the Simpson diversity index (p ≤ 0.05; Figure 7B) in the B group compared to the CK group. For β-diversity, the differences and similarities (Bray-Curtis index) of the maize microbial community between the control group (CK) and the groups treated with the biocontrol bacterium (B) were analysed using Principal Coordinate Analysis (PCoA). As shown in Figure 7C, the samples from the CK and B groups showed distinct clusters.

**Figure 7.**
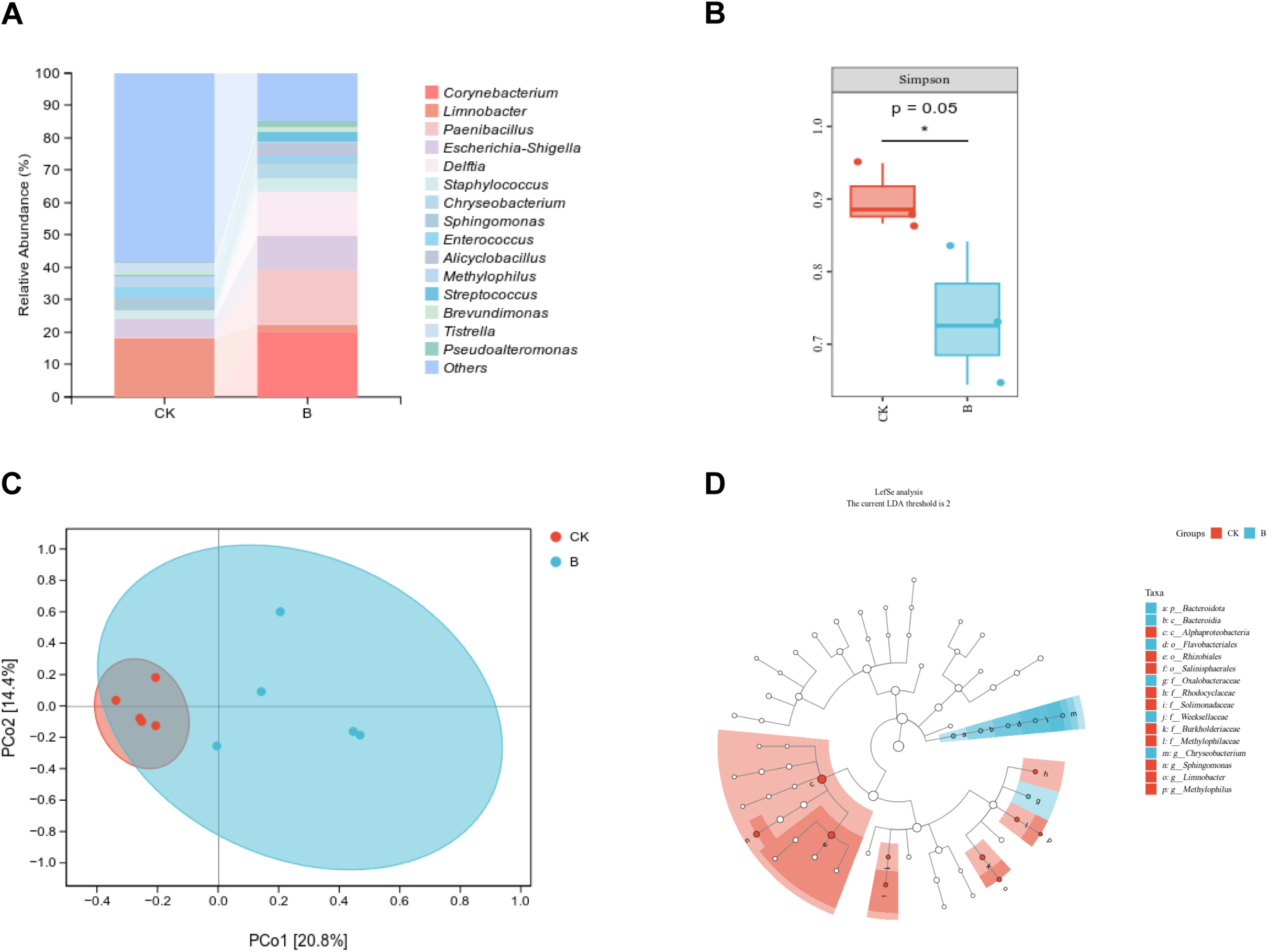
Comparison of microbial community composition between CK and B groups. **(A)** Relative abundance of bacteria community. **(B)** Simpson’s diversity index. **(C)** PCoA score plot. **(D)** LEfSe analysis.

In the CK and B groups, dominant microbial biomarkers were found at the phylum to genus level using linear discriminant analysis effect size (LEfSe). LEfSe difference analysis demonstrated that the abundance of bacterial taxa, such as *Bacteroidota*, *Bacteroidia*, *Flavobacteriales*, *Oxalobacteraceae*, *Weeksellaceae*, and *Chryseobacterium*, were higher in B than in CK, potentially playing major functions in plant disease defense (Figure. 7D). In conclusion, *P*. *peoriae* 3-B4 application resulted in a notable increase of the maize bacterial community diversity, which might contribute to alterations of the community’s function after 3-B4 treatment.

### 3.9 Interactions between microbes and DEGs

To investigate the relationship between microbes and host genes in maize after treatment with biocontrol bacterium *P*. *peoriae* 3-B4, we carried out a genus-level correlation analysis between the top 200 DEGs and 10 microbial taxa.

As Figure 8A illustrates, we observed significant positive correlations (p < 0.05) between most DEGs and the relative abundance of *Delftia*. Additionally, several notable negative correlations (p < 0.05) were observed between the dominant taxa *Limnobacter* and DEGs. The relationships between the microbiota and plant disease resistance-related genes were specifically investigated, and a significant correlation was visually depicted between them through a heat map (p < 0.05; Figure 8B). A statistically significant positive correlation was observed between CCT and *Chryseobacterium* and *Alicyclobacillus*. bZIP had a positive correlation with both *Delftia* and *Chryseobacterium*. Conversely, a notable inverse correlation was observed between HSP and *Limnobacter* and *Sphingomonas*.

**Figure 8.**
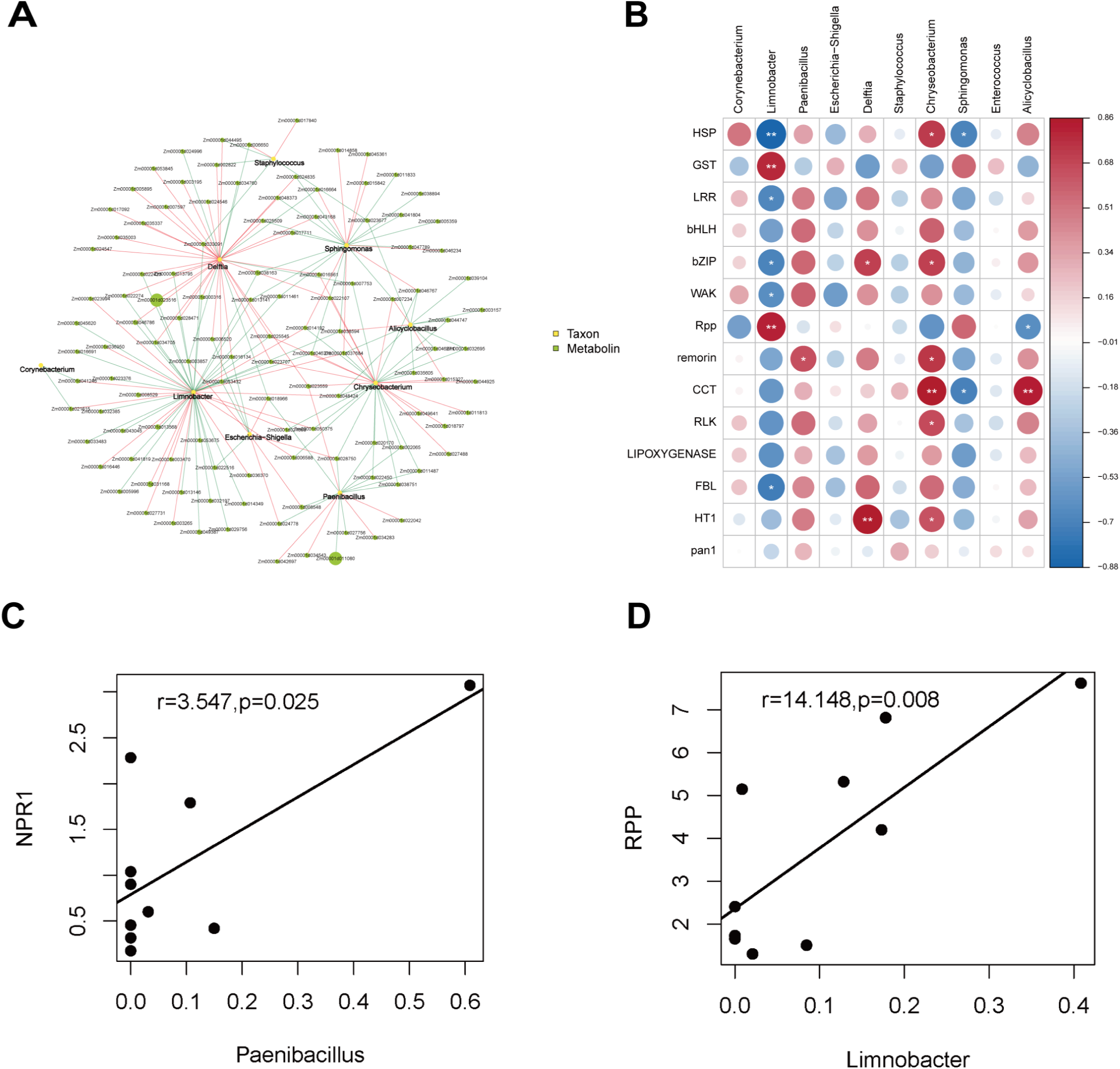
Interactions between the microbes and DEGs related to plant disease resistance. **(A)** Network plot of significant microbe–gene (top 200 DEGs in transcriptome) correlations. Red lines indicate a positive correlation. and green lines a indicate negative correlation. **(B)** Correlation heatmap between microbes and potential genes related to disease resistance. Red indicates a positive correlation, and blue indicates a negative correlation; the color of the correlation indicates the strength of the correlation. The symbol * is used to identify “species–metabolite” pairs with a significant correlation (p<0.05) **(C)** Correlation plots of NPR1 and *Paenibacillus* combinations. **(D)** Correlation plots of RPP and *Limnobacter* combinations.

In addition to the global gene-microbe correlation analyses, Figure 8C, D illustrates the relationships between representative genes and microbial taxa, both of which have previously been linked to host disease resistance. NPR1, an activator of systemic acquired resistance, showed a strong positive correlation with *Paenibacillus* (r=3.547, p=0.025). The NPR1 gene is essential for activating and regulating the plant’s defense mechanisms against a variety of pathogens (38). Furthermore, RPP, an *Arabidopsis* disease resistance gene (43), also exhibited a positive correlation with *Limnobacter* (r =14.148, p=0.008). The proteins encoded by RPP genes are typically members of the nucleotide-binding site-leucine-rich repeat (NBS-LRR) protein family, the function of which is integral to the plant immune system (44,45).

## 4 Discussion

Biocontrol using *in planta* bacteria represents a natural and sustainable method for controlling various plant diseases. Endophytic bacteria, capable of colonizing specific plant cells or tissues and exhibiting resilience to environmental stress, hold significant potential as alternatives to traditional biocontrol agents (46, 47). Microbial fertilizers and biopesticides derived from endophytes and their metabolites are increasingly regarded as promising disease control solutions for future agriculture (47,48).

Previous morphological observations using electron microscopy have revealed that the formation of nodular structures on *F*. *verticillioide*s hyphae during its infection process is primarily associated with infection structures resembling appressorium-like formations at the hyphal tips (49). In contrast to the antifungal effects of *P. peoriae* RP51 reported by Ali et al. (50), where interaction with *Fusarium graminearum* resulted in mycelial shrinkage and distortion, our scanning electron microscopy (SEM) imaging revealed that confrontation with *P. peoriae* 3-B4 induced distinct morphological alterations in *F. verticillioides*, suggesting a different mechanism of action compared to RP51. In research on the antimicrobial properties of *P. peoriae* IBSD35, findings revealed that the pathogen colonies gradually disappeared after the application of the biocontrol bacterium *P*. *peoriae* IBSD35 (51) . These findings indicate that *Paenibacillus* species inhibit pathogens through a range of distinct inhibitory mechanisms.

The *Paenibacillus* genus is well-known for generating antimicrobial compounds such as fusaricidins, pelgipeptin, surfactins, and polymyxins (51,36). Based on the antiSMASH database, *P. peoriae* 3-B4 produces eight antimicrobial metabolites. Paenilan, a conserved ribosomally synthesized peptide found in *Paenibacillus* species worldwide, acts as a novel class I antibiotic with antimicrobial effects against Gram-positive bacteria, including *Bacillus*, *Micrococcus*, and *Paenibacillus* species (52). Tridecaptins, another non-ribosomal antimicrobial peptide synthesized by *Bacillus* and *Paenibacillus* species (53,54), exhibits effective mechanistic activity against Gram-negative bacteria with a low risk of resistance development, potentiating its safe application for environment (55,56). In addition to their antimicrobial properties, fusaricidin B and paeninodin inhibit the growth of phytopathogenic fungi (57,58). Fusaricidin produced by NRPSs has great industrial application potential, given previous findings of its specific inhibitory actions against Gram-positive bacteria and phytopathogenic fungi (59). The antifungal mechanism of fusaricidin involves permeation and disruption of the cell membrane, likely contributing to the broad antifungal spectrum exhibited by *P. peoriae* 3-B4 (7). Although paeninodin is a novel lasso peptide classified under ribosomally synthesized and post-translationally modified peptides (RiPPs), it demonstrates broad antimicrobial and antiviral activity (59).

In the genome of *P. peoriae* 3-B4, we identified 13 genes associated with ISR and PTI, and they exhibited high levels of homology (>95% similarity) with other *P. peoriae* strains, suggesting that *P*. *peoriae* 3-B4 may function in a similar manner to other species within the *Paenibacillus* genus in inducing resistance. Biological control agents can produce systemic resistance inducers, including volatile organic compounds such as methanethiol, isoprene, 2,3-butanediol, and ethyl butyrate, which are released into the environment to counteract pathogen invasion and thus provide protection (31).

After *P*. *peoriae* 3-B4 application, significant changes in the transcriptome of maize occurred, specifically the overexpression of NPR1, bZIP, MYB, LRR, and WRKY. These changes may induce a set of defense mechanisms to combat pathogens directly and indirectly. GST involved in antioxidant defense mechanisms, as it reduces the reactive oxygen species (ROS) levels through its peroxidase activity (60) and thereby protects cells from oxidative damage and enhancing disease resistance (61). Therefore, we hypothesize that the biocontrol effects of *P*. *peoriae* 3-B4 on maize are conferred through inducing specific GST genes and other abovementioned functionally relevant genes.

Moreover, DEGs in the control group and the group treated with the biocontrol bacterium *P*. *peoriae* 3-B4 were mainly enriched in five signaling pathways: plant hormone signaling, MAPK signaling pathway, plant–pathogen interactions, protein processing in the endoplasmic reticulum, and ABC transporter synthesis. These pathways play a crucial role in corn’s defensive response against diseases. The plant hormone signaling and MAPK signal pathways are also induced in tomato plants infected with *Trichoderma afroharzianum* TM24 (62). The results are in line with the findings of Shao et al. (63), in which most DEGs in maize involved in plant hormone signal transduction, MAPK signal pathway, and plant-pathogen interaction pathways. The MAPK signaling pathway plays a crucial part in plant resistance to pathogen infection (64). This pathway involves a cascade reaction of a number of kinases that rapidly respond to external pathogen attack by transducing signals to the nucleus and activating disease resistance genes (65). By regulating multiple defense mechanisms [e.g., disease process-related (PR) proteins synthesis, defense enzymes activation and antimicrobial compounds generation], the MAPK pathway enhances plant defenses against a wide range of pathogens, such as bacteria, fungi, and viruses (66). In addition, the MAPK pathway regulates the transduction of phytohormonal signals (e.g., the ones of jasmonic acid and ethylene), which further orchestrate local and systemic defense responses in plants (67). The MAPK pathway’s downstream components, known as WRKY transcription factors, have the ability to regulate the transcription of genes associated with different defensive reactions in plants. In rice, WRKY67 can induce the expression of genes involved in plant defence, and RNA-Seq analysis has shown that its upregulation significantly enhances resistance to bacterial blight (68). The genes involving the MAPK pathway were also induced in our study, with their increased expression potentially contributing to enhanced pathogen resistance in maize.

Microbiota is widely recognized for its functions in regulating plant immunity, promoting plant growth, and maintaining overall host health. Current research into the application of biocontrol agents has mainly centred on the alteration of community composition of phyllosphere microbiota, rhizosphere microbiota, and in the soil. However, the in planta microbiota has been relatively little studied. Jia et al. (69) and Wang et al. (70) found that the microbial community structure and the relative abundance of some beneficial genera significantly changed in the biocontrol treatment groups compared with CK in the phyllosphere and soil, indicating that BCA can enhance the composition of microbial communities. In this study, Simpson’s diversity index was much higher in the CK group than the *P*. *peoriae* 3-B4 treatment group. Additionally, the treatment group exhibited higher levels of beneficial microbiota, including *Paenibacillus*, *Delftia*, *Chryseobacterium*, and *Pseudomonas*. Of these, *Paenibacillus* is commonly found in plants and performs nitrogen fixation, promoting plant growth and biocontrol properties. *Delftia* can degrade various pollutants, acts as a biological control, and promotes plant growth through multiple mechanisms, such as producing siderophores and antimicrobial compounds (71). *Chryseobacterium* contributes to the health of plant root systems through nutrient cycling, delivering plant growth-regulating hormones, and exhibiting biocontrol capabilities against plant pathogens (72). Furthermore, in combination with the diversity analysis results, we concluded that *P*. *peoriae* 3-B4 treatment increases microbiota diversity, especially beneficial microorganisms, in maize. These results collectively indicate that biocontrol agent treatment not only alters the microbial community composition in the phyllosphere, rhizosphere, and soil but also affects the internal microbiome of the host plant. In turn, this influences the composition and role of microbial populations across different plant niches, thereby playing a role in pathogen resistance.

In our study, multi-omics analysis was employed to understand the efficacy of *P*. *peoriae* 3-B4 application on *F*. *verticillioides* (Figure 9). Based on the *P*. *peoriae* 3-B4 genome, genes that induce resistance were identified. When activated, these genes trigger the plant’s immune response, including plant hormone signaling, ROS burst, activation of MAPK and defense genes, shifts in transcriptional regulation, and the synthesis of secondary metabolites (73,74,75,76). Transcriptome analysis indicated a significant upregulation of pathways and gene expression levels associated with disease resistance in maize following biocontrol bacterium *P*. *peoriae* 3-B4 application. Through transcriptome data screening, NPR1, a key regulatory protein that is vital for plant defense, was identified as significantly upregulated. Additionally, 16S rRNA analysis demonstrated that *P*. *peoriae* 3-B4 treatment not only resulted in significant alterations to the microbial community composition, with a higher Simpson’s index than in the control group, but also led to an increase in beneficial microbial community diversity, including *Paenibacillus*, *Delftia*, and *Corynebacterium*. Further transcriptome and microbial diversity association analyses revealed a positive correlation between NPR1 expression and beneficial microbes, especially *Paenibacillus* species. This interaction highlights the important function of beneficial microbes in enhancing plant immune responses. Overall, the combined activation of resistance synthesis genes, upregulation of NPR1, and activation of plant immunity, including SA accumulation and defense gene expression show that *P*. *peoriae* 3-B4 may induce the transcription levels of disease resistance pathways and synergistically affect the composition and function of the microbiome to enhance host plant’s immunity against pathogens, ultimately improving its overall health.

**Figure 9.**
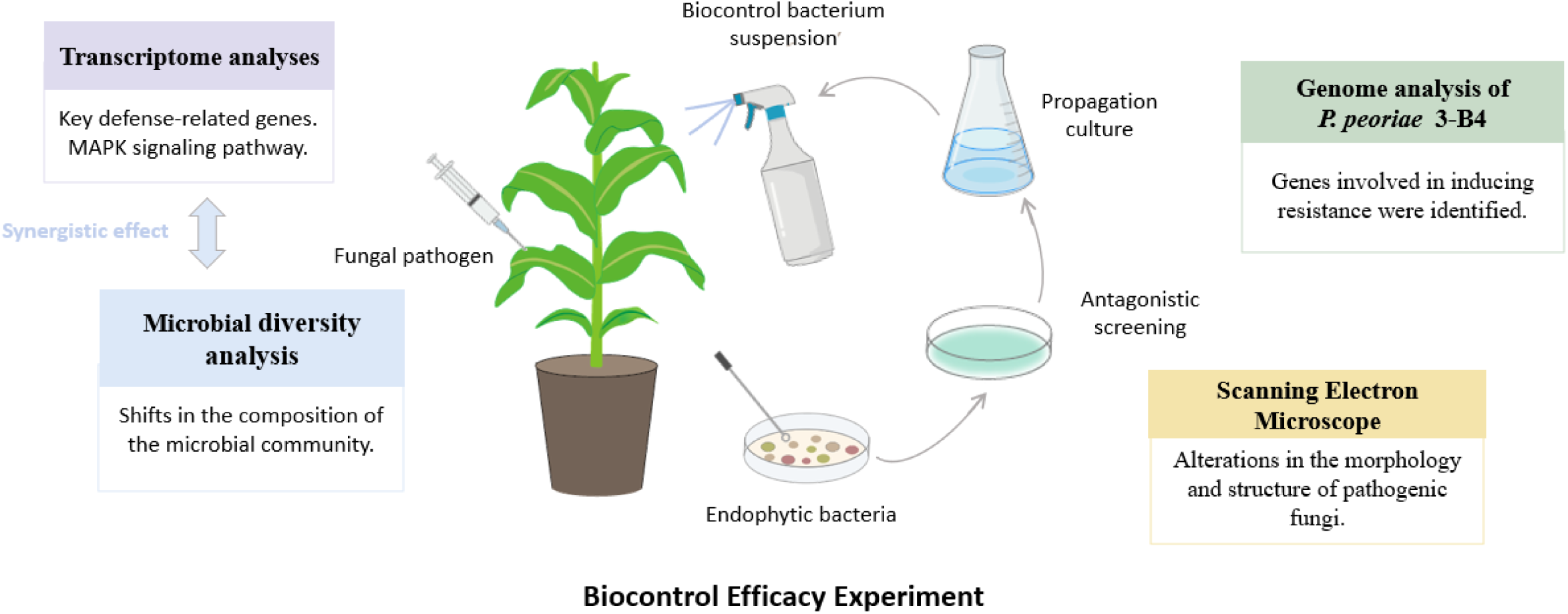
Mechanisms of biocontrol bacterium *Paenibacillus peoriae* 3-B4 in enhancing maize disease resistance.

## Conclusion

In this study, we isolated and characterized a potent biocontrol strain, *P*. *peoriae* 3-B4, which enhanced maize resistance to *F*. *verticillioides*. In the genome of *P. peoriae* 3-B4, genes related to secondary metabolite production and resistance induction, including 8 gene clusters and 13 genes linked to ISR and PTI, were mined and analyzed. Transcriptomic results revealed that the DEGs were primarily associated with 5 signaling pathways and that some critical genes, such as NPR1, bZIP, MYB, LRR, and WRKY, were regulated. Furthermore, 16S rRNA analysis showed that biocontrol bacterium *P*. *peoriae* 3-B4 application altered the bacterial community composition, and some beneficial microbial communities showed a positive correlation with DEGs in the transcriptome. Overall, our study provides evidence that *P*. *peoriae* 3-B4 application can improve disease resistance. This finding may enable further investigation of the mechanisms underlying gene expression levels and microbial structural changes by which biocontrol bacteria enhance plant’s resistance to pathogens. Additionally, our study revealed that disease resistance was enhanced during the *P*. *peoriae* 3-B4 induction process by co-regulating disease resistance genes and functional microbial abundance. By revealing the mode of action of a natural maize endophyte in managing maize seedling blight, this work offers a foundation for development of novel biocontrol agents. These agents will not only reduce the reliance on chemical pesticides, thereby minimizing environmental pollution and enhancing food safety, but also contribute to sustainable agriculture, food security, and improved agricultural product quality.

## Author Statement

The authors declare that they have no known competing financial interests or personal relationships that could have appeared to influence the work reported in this paper.

## CRediT authorship contribution statement

Yue Hu: Writing - Original draft preparation, Formal analysis, Investigation. Yifan Chen: Methodology, Conceptualization. Software. Shengqian Chao: Visualization, Data Curation. Yin Zhang: Project administration. Lili Song: Data curation, Validation. Hui Wang: Software, Validation, Project administration. Yingxiong Hu: Resources. Beibei Lü: Supervision, Conceptualization, Funding acquisition.

